# Zymosan-induced leukocyte and cytokine changes in pigs: a new model for streamlined drug testing against severe COVID-19

**DOI:** 10.1101/2022.09.23.509252

**Authors:** Gábor Kökény, Tamás Bakos, Bálint András Barta, Georgina Viktória Nagy, Tamás Mészáros, Gergely T. Kozma, András Szabó, János Szebeni, Béla Merkely, Tamás Radovits

## Abstract

Injection of 0.1 mg/kg zymosan in pigs i.v. elicited transient hemodynamic disturbance within minutes, without major blood cell changes. In contrast, infusion of 1 mg/kg zymosan triggered maximal pulmonary hypertension with tachycardia, lasting for 30 min. This change was followed by a transient granulopenia with a trough at 1 h, and then, up to about 6 h, a major granulocytosis, resulting in a 3-4-fold increase of neutrophil-to-lymphocyte ratio (NLR). In parallel with the changes in WBC differential, qRT-PCR and ELISA analyses showed increased transcription and/or release of inflammatory cytokines and chemokines into blood, including IL-6, TNF-α, CCL-2, CXCL-10, and IL-1RA. The expression of IL-6 peaked at already 1.5-2.5 h, and we observed significant correlation between lymphopenia and IL-6 gene expression. While these changes are consistent with zymosan’s known stimulatory effect on both the humoral and cellular arms of the innate immune system, what gives novel clinical relevance to the co-manifestation of above hemodynamic, hematological, and immune changes is that they represent independent bad prognostic indicators in terminal COVID-19 and other diseases involving cytokine storm. Thus, within a 6 h experiment, the model enables consecutive reproduction of a symptom triad that is characteristic of late-stage COVID-19. Given the limitations of modeling cytokine storm in animals and effectively treating severe COVID-19, the presented relatively simple large animal model may advance the R&D of drugs against these conditions. One of these disease markers (NLR), obtained from a routine laboratory endpoint (WBC differential), may also enable streamlining the model for high throughput drug screening against innate immune overstimulation.

## Introduction

Cytokine storm syndrome (CSS), a hyper-inflammation characterized by abnormally high levels of proinflammatory cytokines and other immune mediators in blood, is known to be a major contributor to the death of patients with severe COVID-19.^1-17^ Among the cytokines analyzed in the present study, high serum IL-6 and IL-1RA levels were found as independent risk factors for mortality from COVID-19.^18-23^ CSS is, however, not the only sign of bad prognosis in late-stage COVID-19. Another one is the association of neutrophilia with lymphopenia, manifested in a rise of neutrophil/lymphocyte ratio (NLR),^24,25^ and yet another is hemodynamic derangement, such as pulmonary hypertension^26-28^ and/or systemic hypotension.^29,30^

Peculiarly, hemodynamic derangement and WBC differential changes are also characteristic features of the pigs’ response to i.v. administered nanoparticles. The experimental setup, referred to as porcine complement (C) activation-related pseudoallergy (CARPA) model,^31-35^ has been used for the screening of nanoparticulate drugs (nanomedicines) for potential immune reactivity, manifested in infusion reactions. The model utilizes zymosan, a strong activator of the innate immune system, as a highly reproducible positive control, since zymosan’s robust pulmonary hypertensive and systemic hypotensive effects are observed essentially in all pigs.^32-34^ These effects, taken together with the fact that zymosan is known to induce proinflammatory cytokines in murine models^36,37^ and human peripheral blood mononuclear cells (PBMCs)^38^ led to the hypothesis that zymosan could be used to induce in pigs the above triad of stigmatic endpoints of severe COVID-19.

Accordingly, the goal of the present study was to examine if the adverse hemodynamic and hematological effects of zymosan could be associated with inflammatory cytokine release in pigs, and if yes, to propose a simplified model of late-stage COVID-19 that may allow streamlined drug testing against this, and other diseases involving CSS.

## Materials and Methods

### Materials

Ficoll-Paque was obtained from GE Healthcare Bio-Sciences AB (Uppsala, Sweden). The porcine C3a kit was obtained from TECO*Medical* AG, Sissach, Switzerland (Cat No: TE1078). Zymosan, Dulbecco’s phosphate-buffered saline (PBS) without Ca^2+^/Mg^2^ were from Sigma Chemical Co. (St. Louis, MO, USA). Pre-designed primers for IP10 (CXCL10, Assay cID: qSscCED0019399) and IL6 (Assay ID: qSscCED0014488) were purchased from BioRad laboratories (Herkules, CA).

### Animals

Landrace pigs were obtained from the Research Institute for Animal Breeding, Nutrition and Meat Science of the Hungarian University of Agriculture and Life Sciences (Herceghalom, Hungary). The study involved 13 female and castrated male pigs in the 22-32 kg size range. The experiments were approved by the Ethical Committee of Hungary for Animal Experimentation (permission numbers: PE/EA/843-7/2020 and conformed to the EU Directive 2010/63/EU and the Guide for the Care and Use of Laboratory Animals used by the US National Institutes of Health (NIH Publication No.85–23, revised 1996).

### Anesthesia and instrumentation

Animals were sedated with an im. injection of 25 mg/kg ketamine (50mg/mL, Gedeon Richter Plc. Budapest, Hungary) and 0.3 mg/kg midazolam (15mg/3mL, Kalceks AS, Riga, Latvia) and were carefully transported into the laboratory. Anesthesia was induced with a propofol bolus through an auricular vein. Airways were secured by insertion of an endotracheal tube. Unless otherwise specified, animals were allowed to breathe spontaneously during the experiments. Controlled ventilation was applied in case of animals, where continuous measurement of hemodynamic parameters necessitated invasive surgical interventions. Surgery was done after povidone iodine (10%) disinfection of the skin. To measure the PAP, a Swan-Ganz catheter (AI-07124, 5 Fr. 110 cm, Arrow International Inc.) was introduced into the pulmonary artery via the right internal jugular vein. A Millar catheter (SPC-561, 6 Fr. Millar Instruments, Houston, TX, USA) was placed into the left femoral artery to record the systemic arterial pressure (SAP). Additional catheters were introduced into the left external jugular vein for drug administration, into the left femoral vein for venous blood sampling and the right common carotid artery for arterial blood gas analysis.

The latter was executed with a Roche COBAS B221 benchtop analyzer. Hemodynamic and ECG data were collected using instruments from Pulsion Medical Systems, and Powerlab, AD-Instruments (Castle Hill, Australia). Furthermore, end-tidal pCO2, ventilation rate and body temperature were also continuously measured, but they served no information beyond the measurements presented in this study and are therefore not shown.

### Experimental protocols in different stages

This study was performed in 3 stages to measure the acute (minutes) and subacute (hours to days) effects of zymosan in animals dedicated only to this study and in others, where zymosan was used as control. In the latter case we used animals where other treatments caused no, or minimal physiological changes, and it could be ascertained that the other treatments had no impact on zymosan’s effects. In the first stage 5 pigs were treated with bolus injection of 0.1 mg/kg zymosan, and the resultant hemodynamic, hematological and blood immune mediator changes were monitored or measured as described below. In the second stage 1 mg/kg zymosan was infused in 4 pigs over 30 min, and the same protocol was applied as in stage 1, except that the monitoring and serial blood sampling lasted for 6.5 h. In the third stage 5 pigs were infused with 1 mg/kg zymosan, followed by blood withdrawals at 10, 20, 30, min, then in increasing times until 6.5 h, and then at 2-3 days intervals for 15 days.

### Blood Assays

As described above, 10 mL of venous blood was drawn from the pigs at different times into EDTA containing vacuum blood collection tubes (K_3_EDTA, Vacuette). 0.5 mL of blood was aliquoted for use in an ABACUS Junior Vet hematology analyzer (Diatron, Budapest, Hungary) to measure the following parameters of blood cells: white blood cell (WBC), granulocyte (GR) and lymphocyte (LY), platelet (PLT), red blood cell (RBC) count and hemoglobin (Hgb) concentration. For measuring TXB_2_, a stable metabolite of TXA_2_, 4 μg indomethacin (diluted in 2 ul of 96% ethanol) was mixed to 2 mL of anticoagulated blood to prevent TXA_2_ release from WBC before centrifugation at 2000g, for 4 min at 4°C. Another 2 mL of anticoagulated blood was directly centrifuged using the same settings to separate the plasma. After centrifugation the plasma samples were aliquoted, frozen, and stored at -70°C until the TXB_2_ assay, as described in the kits’ instructions, and the cytokine assays, as described below.

### Cytokine Measurements

Cytokine levels were measured in plasma samples derived from blood, as described above. which were taken in the previously discussed timepoints during and after zymosan infusion. The levels of IL-1β, IL-6, IL-8 and TNFα cytokines were determined using a High Sensitivity 4-Plex Porcine Cytokine kit from Quansys Biosciences Inc. (West Logan, UT, USA) in accordance with the protocol provided by the manufacturer. The data was collected with Imager LS by Quansys operated through Q-View Software, which was also used to evaluate the results.

### Isolation of PBMC and quantitative RT-PCR

Peripheral blood mononuclear cells (PBMC) were isolated from 4 ml anticoagulated blood within 30 minutes at each experimental time point. Briefly, 2 ml blood was transferred into a 15 ml tube and diluted with 2 ml phosphate-buffer saline (PBS pH 7.4). In a new 15 ml tube, 3 ml Ficoll-Paque media (GE Healthcare) was pipetted into the bottom and the diluted 4 ml blood sample was carefully layered on top, centrifuged for 30 min at 400g. The upper plasma layer was removed, and the leukocyte layer was transferred into a new tube containing 6 ml PBS, washed and centrifuged. The PBMC pellet was resuspended in 1 ml TriZol (Invitrogen) and total RNA was extracted according to manufacturer’s instructions. RNA pellet was resuspended in RNAse-free water and the RNA concentration was determined photometrically on a NanoDrop microphotometer (Thermo). One microgram of RNA of each sample was reverse transcribed with the High-Capacity cDNA Reverse Transcription kit from Applied Biosystems (Applied Biosystems/Life Technologies, Carlsbad, CA, USA) using random primers in a final volume of 20 μl. Quantitative real-time PCR reactions were performed on a Bio-Rad CFX96 thermal cycler (Bio-Rad Hungary, Budapest, Hungary) using the SensiFast SYBR Green PCR Master Mix (Thermo Fisher). The specificity and efficiency of each PCR reaction was confirmed with melting curve and standard curve analysis, respectively. Each sample was quantified in duplicates and normalized to 18S rRNA *(RN18S)* expression. Mean expression values were calculated as fold expression relative to a baseline control sample using the *2*^*-*ΔΔ*Ct*^ formula. Pre-designed primers for IP10 (CXCL10, Assay ID: qSscCED0019399) and IL6 (Assay ID: qSscCED0014488) were purchased from BioRad laboratories. Primer sequences for RN18S, IL1RA and CCL2 are shown in Table 1.

**Table 1.**
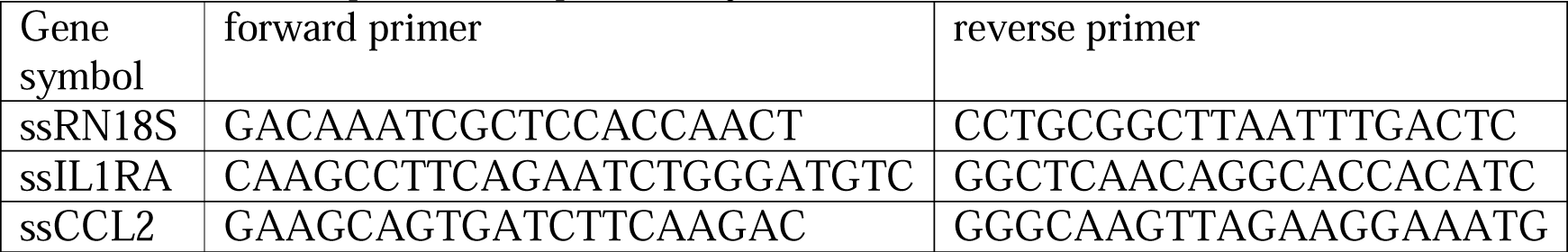
Primer sequences for qPCR analysis

### Statistical analyses

All data are presented as mean ± SD. Statistical analysis was performed using SPSS 10 (IBM, USA). PBMC gene expression values were evaluated Kruskal-Wallis test and Dunn’s post-hoc test. Blood cell counts and serum cytokine protein levels were evaluated with ANOVA followed by Dunnett’s multiple comparison test. Level of significance was set to p<0.05 in each analysis.

## Results

### Early hemodynamic, hematological, and immune mediator changes caused by low-dose bolus injection of zymosan

As first step in pursuing the hypothesis delineated in the introduction, we reproduced the robust hemodynamic changes caused by a single bolus injection of 0.1 mg zymosan in pigs. As shown in Figure 1, the cardiopulmonary reaction starts with a sudden rise of PAP (A), fall of SAP (C) and massive release of TXB2 (G) exactly paralleling the PAP. The heart rate (HR, E) and blood cells (B, D, F). showed no major changes although a small, statistically significant decline of granulocyte count was detectable (D). The SAP returned to baseline within 10 min, while for PAP and HR it took longer time (up to 30 min, not shown) to return to near normal levels.

**Figure 1.**
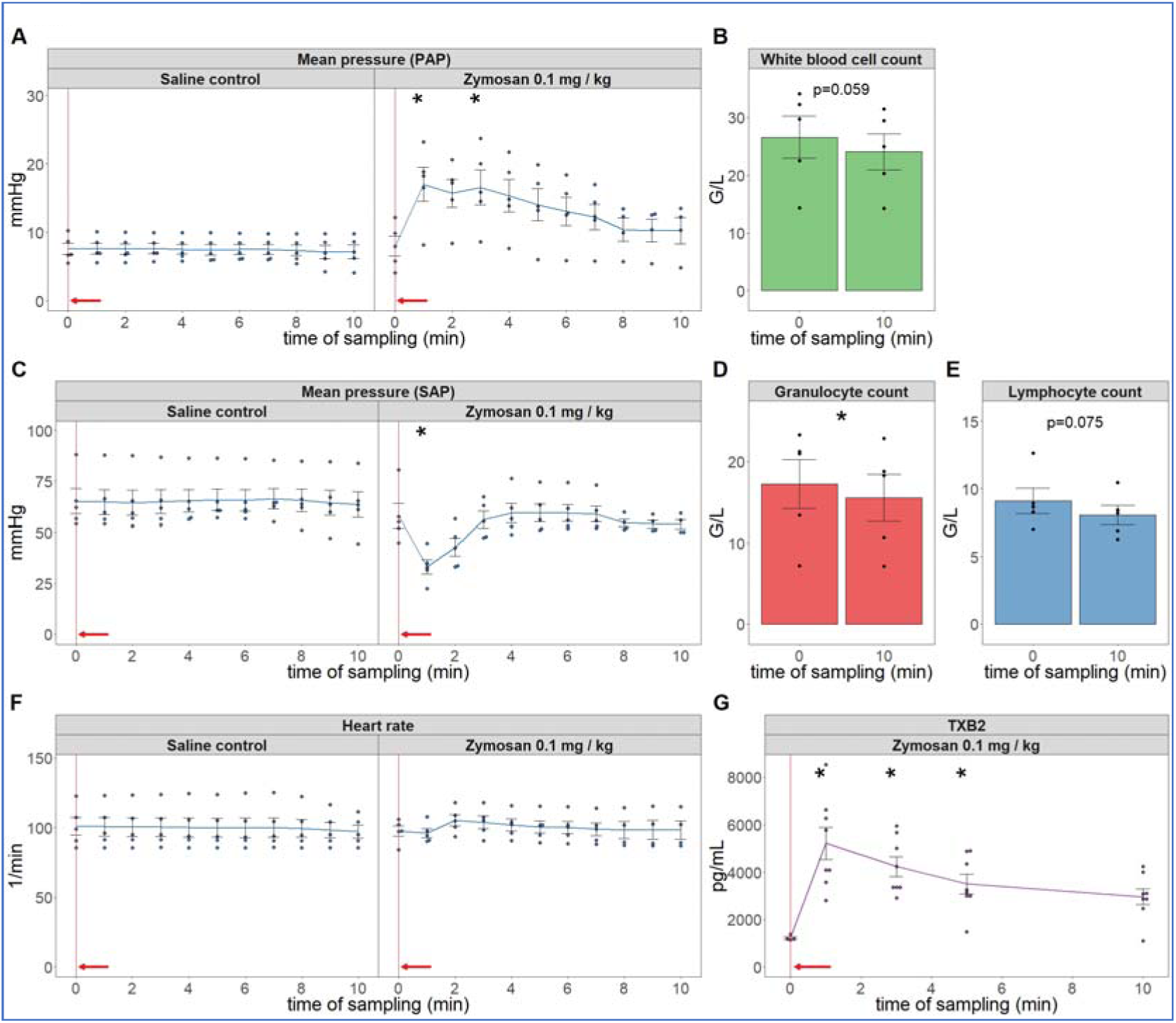
Physiological changes caused by bolus injection of 0.1 mg/kg zymosan in pigs. In A, C, E, the PAP, SAP and HR were continuously recorded, and the coinciding values were averaged (±SEM) in 4 animals every minute over 10 min both before and after the zymosan injection. Blood white blood cell count (B), granulocytes (D) and lymphocytes (F) were counted in a coulter counter at 10 min after zymosan injection and were related to their respective baseline, i.e., the last blood collection before the zymosan injection (0 min). TXB2 increased 4-fold within a minute after zymosan administration (G). G/L means cell number *10^6^/liter, * p<0.05 relative to respective baselines, determined by paired t-test.

### Extended follow-up of hemodynamic, hematological, and immune mediator changes caused by high dose zymosan infusion

In lack of major neutrophilia, the mentioned symptom of late-stage COVID-19 that we were trying to identify, we increased the administered dose of zymosan 10-fold and gave it in infusion, rather than bolus. Because the hemodynamic monitoring is invasive, this experiment had to be terminated after 6.5 observation period. Figure 2 shows the hemodynamic changes after initiation of the zymosan infusion. Interestingly, the infusion was associated with maximal rise of PAP, but after completing it, the pulmonary pressure normalized within 20 min. The SAP showed major fall during infusion and then slow return to normal in 1 of 4 animals; a measure that has proven to be very variable in all previous CARPA studies.^39-44^ The heart rate and exhaled CO_2_, an indicator of pulmonary function, showed no significant differences. These data imply an immediate cardiovascular effect of zymosan, which can be explained with immediate TXA_2_ release with entailing pulmonary vasoconstriction. Of particular importance regarding the hypothesis of this study, the blood cell changes did show the expected granulocytosis with lymphopenia, between about 1 and 6 h after starting the 30 min infusion. This effect is shown in Fig 3, together with data from 5 more pigs, which were not subjected to invasive blood pressure recording and, thus, were not sacrificed after 6.5 h. These animals were subjected to blood withdrawals for blood cell counting and cytokine analysis up to 15 days, whose results are presented below.

**Figure 2.**
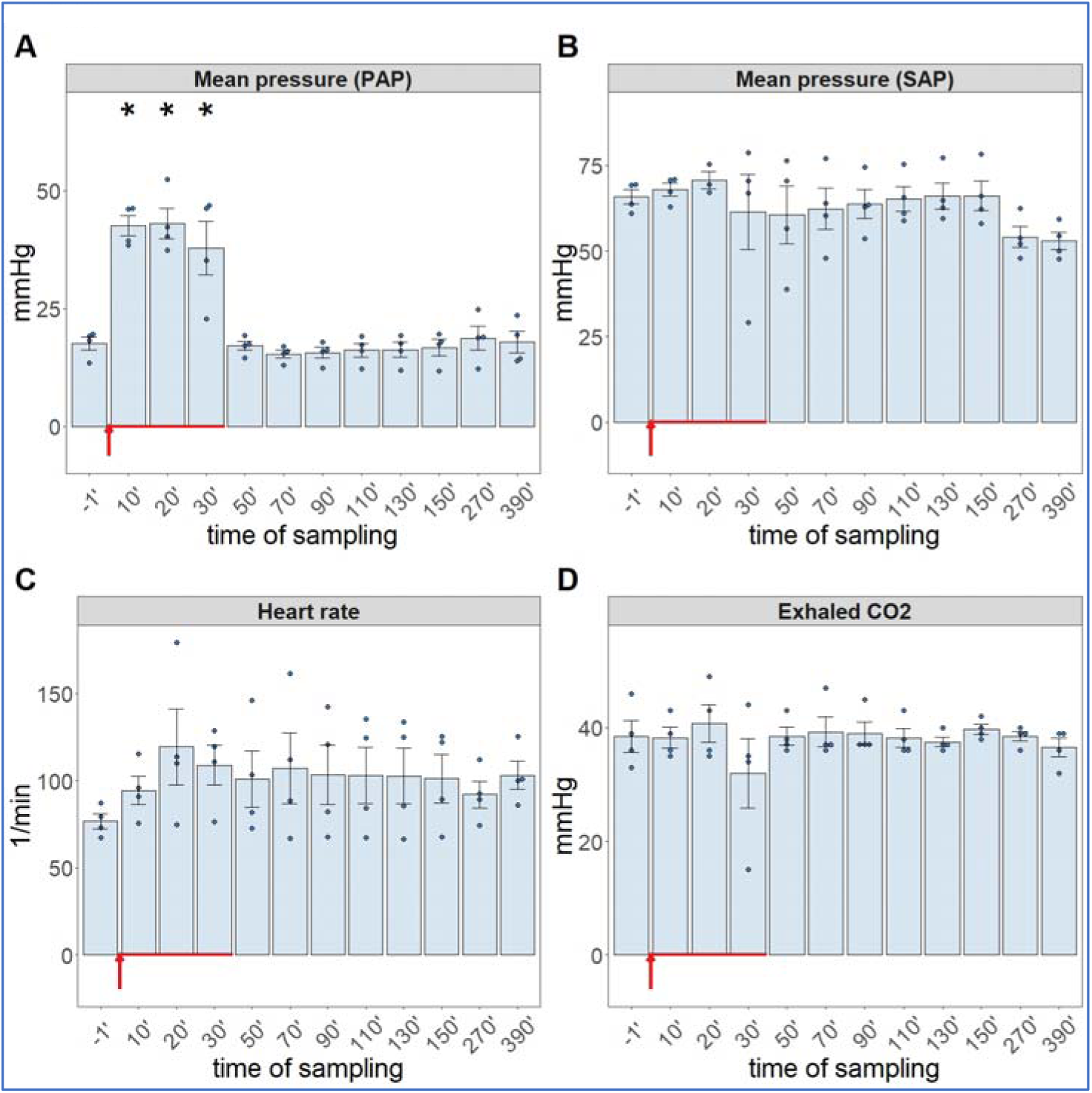
Physiological changes caused by infusion of 1.0 mg/kg zymosan in pigs. PAP (A), SAP (B), HR (C) and exhaled CO_2_ (D) were continuously recorded up to 6.5 h, and the coinciding values at the indicated time points were averaged (±SEM) in 4 animals. Here we show the blood pressure on absolute (mmHg) scale without the flat pre-injection background (Fig 1). The red arrow indicates zymosan administration.

**Figure 3.**
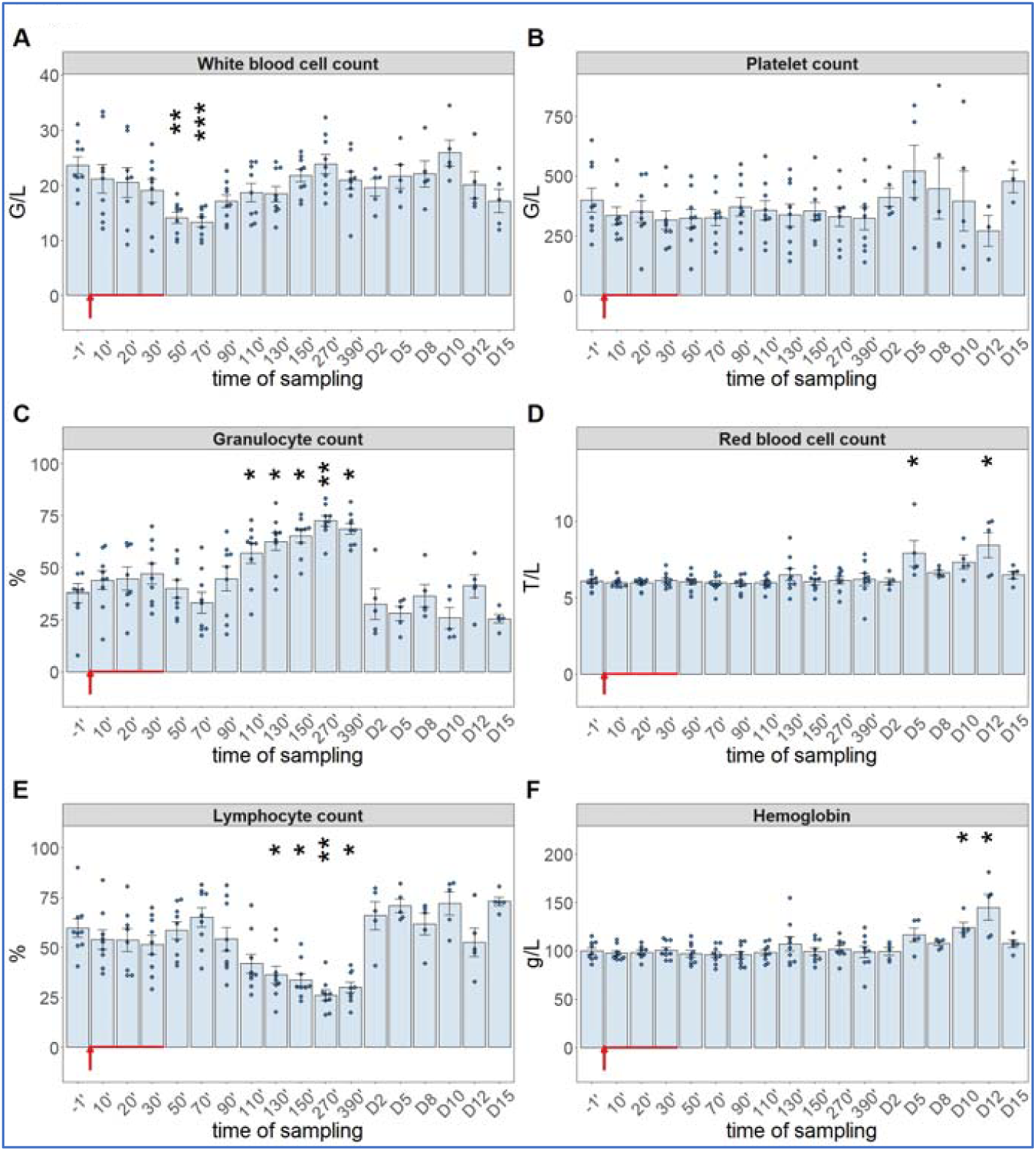
White blood cell count (A), platelet count (B), granulocyte % (C), RBC count (D), lymphocyte % (E), hemoglobin levels at different sampling times. Until 390 min, the data have been merged from the 2nd (n = 4 pigs) and 3rd phase (n = 5 pigs) of the study, and their overlapping exclude a pro-inflammatory influence of hemodynamic monitoring (surgery). ANOVA test, *p<0.05 vs 0’; ** p<0.01 vs 0’; *** p<0.001 vs 0’.

### Long-term follow-up of hematological, TXB2 and cytokine changes caused by high dose zymosan infusion

As shown in Fig. 3, we observed significant drop in WBC count around 1 h after the start of zymosan infusion (Fig 3A), which did not involve changes in neutrophil granulocyte/lymphocyte ratio (NLR, about 4/6) (Fig 3 C, E). However, thenceforwards, while the WBC count returned to normal in about 3-4 h (Fig 3A), the NLR gradually rose up to 7/3, near 3-4-fold relative to the baseline ratio. In absence of absolute (total) increase or decrease in WBC count at the peak of NLR (Fig 3A, C, E), the above changes imply a relative leukocytosis with absolute lymphopenia, a shift in WBC differential in favor of the innate, nonspecific versus the acquired, specific antimicrobial immune response. The platelet count did not change over time (Fig. 3B), and we did not measure consistent trends in RBC counts or blood hemoglobin levels, either, although on some days the numbers showed significant rises relative to baseline (Fig. 3D, F).

Regarding the cytokine changes, the PCR assay showed significant expression of CCL-2 (Fig 4A), CXCL10 (B), IL-1RA (C) and IL-6 mRNAs in close parallelism with the changes in WBC differential, with peaks in the 90-270 min range. The relative mRNA expressions decreased in the above order, while in terms of speed, IL-6 mRNA had faster rise than those of the other 3 cytokines (peaking at already 90-110 min vs. 130-150 min). We also detected significant increases in IL-6 gene transcription in 2 of 5 animals on day 8 and 10, again (Fig. 4D).

**Figure 4.**
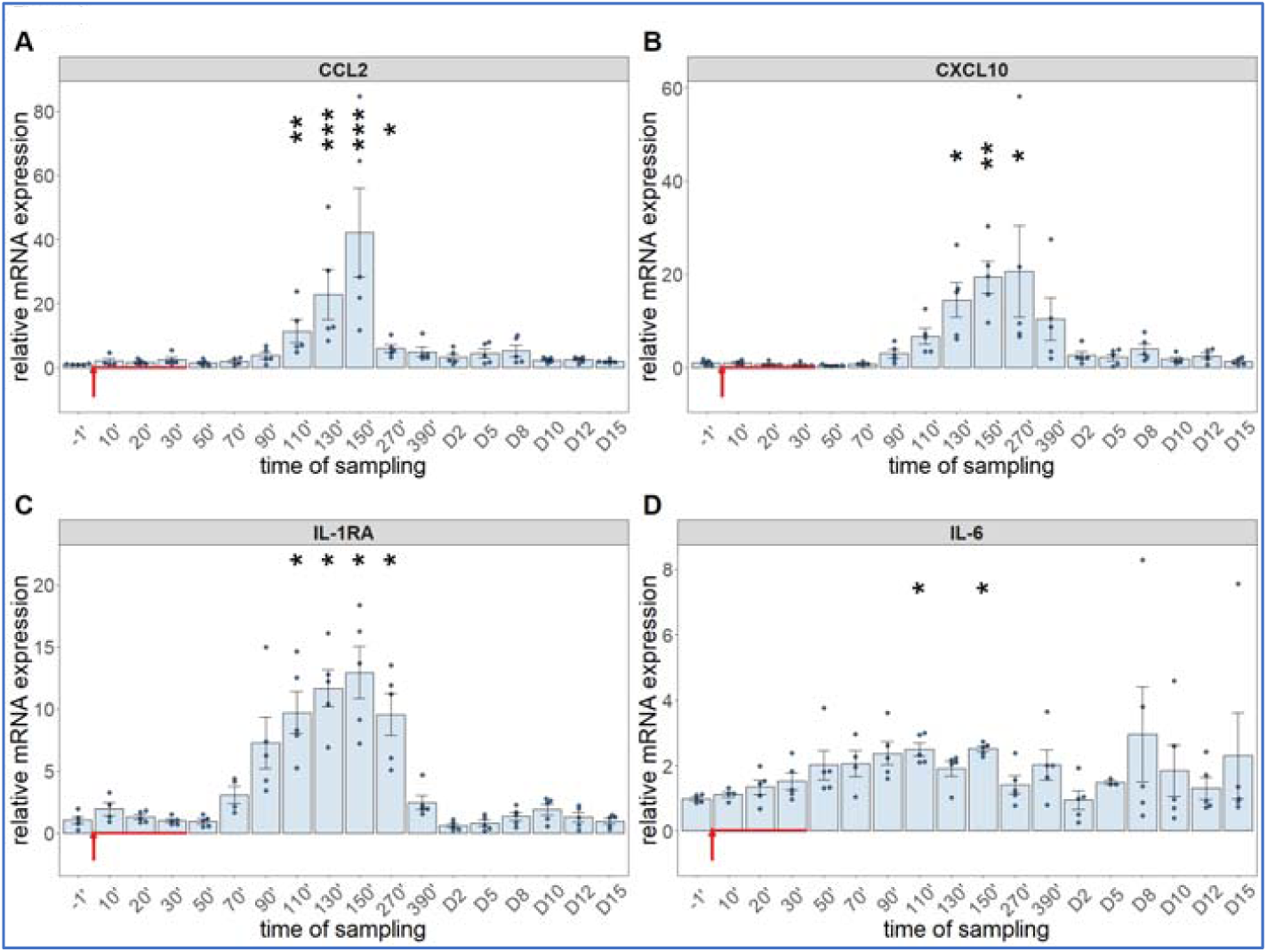
Gene expression of CCL2 (A), CXCL10 (B), IL-1RA (C) and IL-6 (D) were assessed from PBMCs isolated at different sampling times. Kruskal-Wallis test, *p<0.05 vs 0’; ** p<0.01 vs 0’; *** p<0.001 vs 0’

The cytokine protein assays performed for IL-6 and others (whose mRNA expression were not analyzed, i.e., TNF-α, IL-8 and IL-1β) confirmed the early expression of IL-6, but showed the production of TNF-α to be even faster and more intense, peaking about 30 min earlier at about 10-fold higher concentration than IL-6 (Fig 5A and B). The concentrations of IL-8 and IL-1β tended to be increased in some animals after day 10, with substantial individual variation (Fig 5C, D).

**Figure 5.**
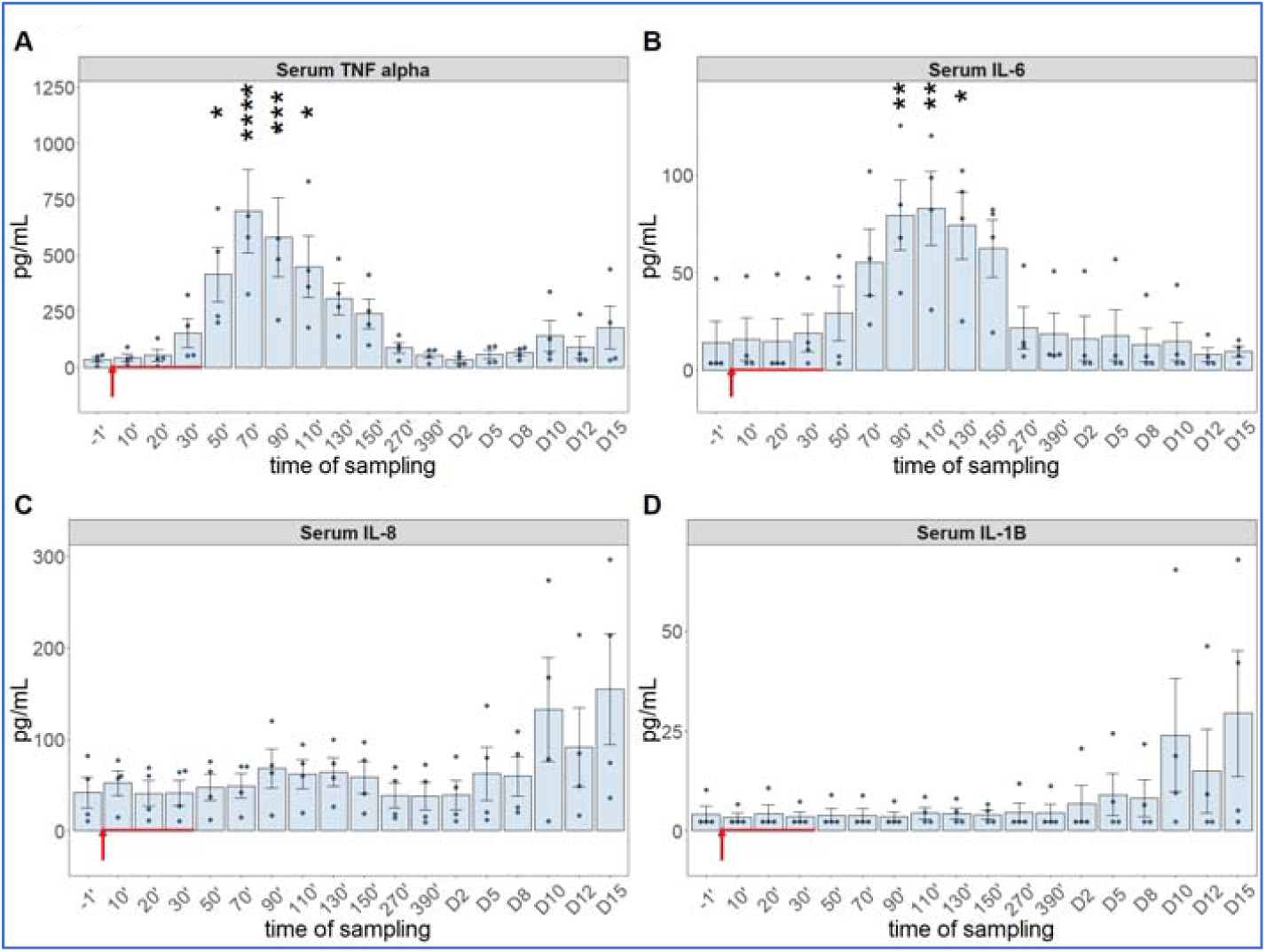
Serum cytokine levels at different sampling times (ELISA). ANOVA, with Dunnett’s multiple comparison test, *p<0.05 vs 0’; ** p<0.01 vs 0’; *** p<0.001 vs 0’; **** p<0.0001 vs 0’

Comparing the kinetics of NLR changes with cytokine transcription or translation revealed overlays with all cytokines analyzed (Fig. 6A-F), except IL-8 and IL-1β, which did not show changes in the 30-270 min period (Fig. 5C, D). Fig. 6A-F also shows that the rise of NLR, which reflects a shift in cellular immune response towards innate defense, started after about 1 h and peaked at 5h (blue line in Fig 6A-F). The cytokines whose early rises and peaks consistently preceded these changes of NLR were TNF-α and IL-6 (Fig. 6A-C). In fact, for IL-6, we could show highly significant correlation between lymphopenia and IL-6 mRNA expression (Figure 7). Nevertheless, it would be premature to point to certain cytokines versus others as sole, or most significant contributors to the blood cell changes, since there were also overlaps between the changes of NLR and expression CCL2, IL-1RA and CXCL-10 genes (Fig 6 D-F). Notably, we do not know the responsiveness of blood cell changes to different cytokines. Thus, what can certainly be stated is that cytokine production correlated with the rise of NLR, which is consistent with the coincidence of NLR rise and CSS in severe COVID-19.

**Figure 6.**
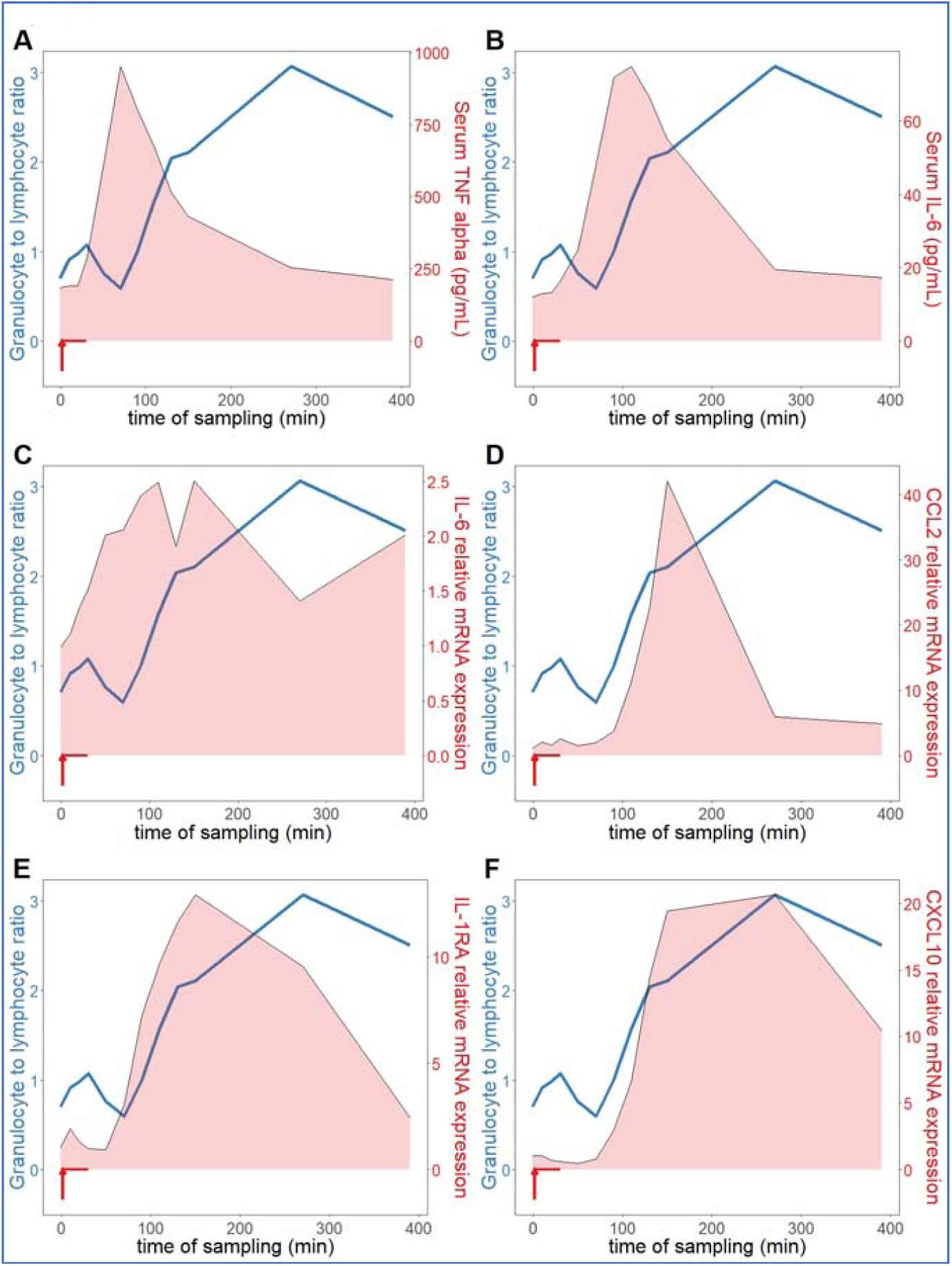
Relationship between serum cytokine levels and NLR. The NLR (neutrophil granulocyte-to-lymphocyte ratio (blue line), increased continuously in an hour after zymosan administration, which was preceded by increased TNF-α (A) and IL-6 (B, C). The expression of CCL2 (D), IL-1RA (E) and CXCL10 (IP-10, F) genes proceeded in parallel with the rise of NLR (n=9).

**Figure 7.**
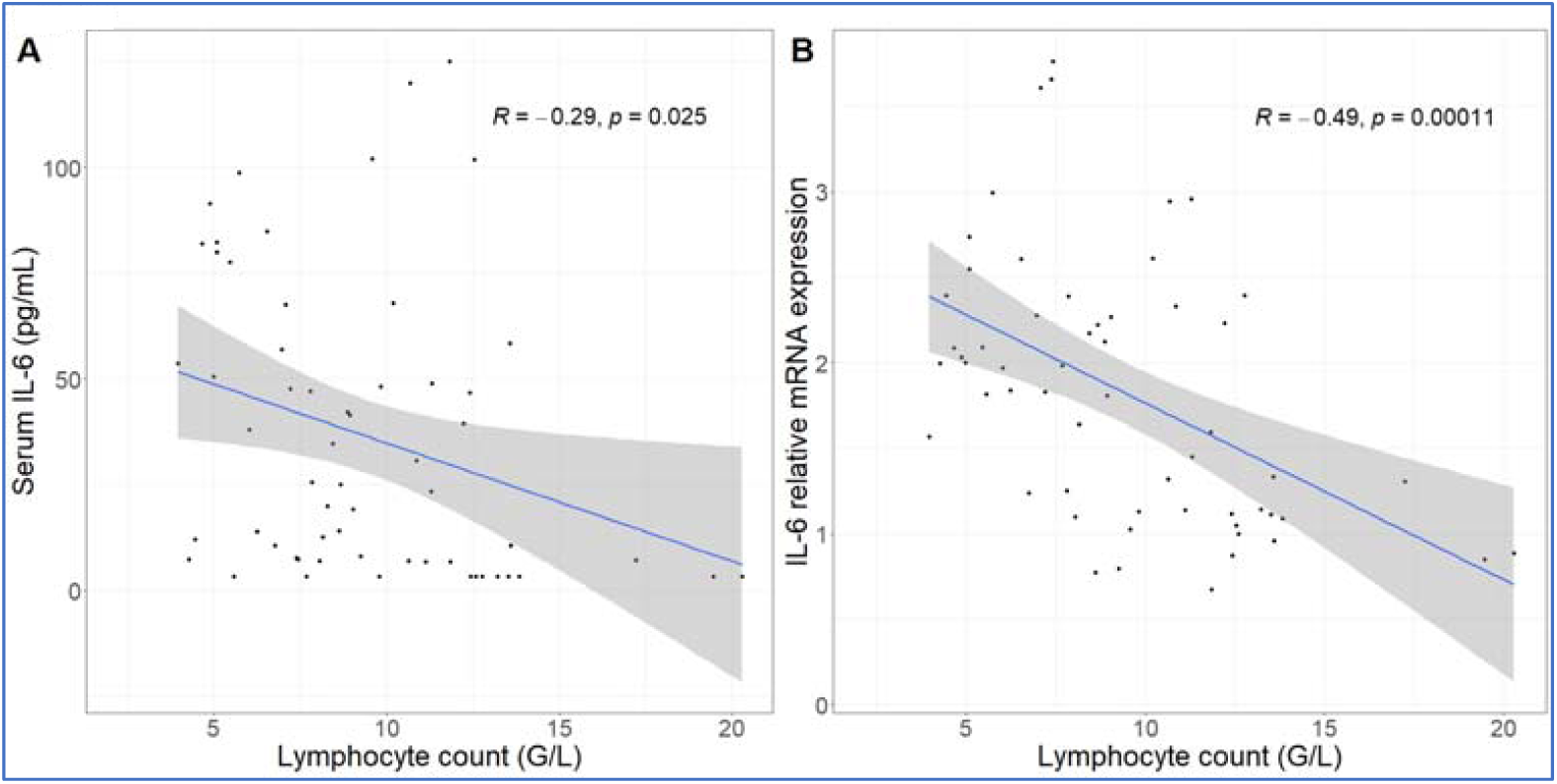
Correlation between IL-6 protein secretion and gene expression and absolute lymphocyte counts. The lymphopenia significantly correlated with increased IL-6 protein (A) and mRNA expression (B) over time.

## Discussion

### The structure and immune effects of zymosan

To help understand how a short infusion of yeast microparticles (zymosan) can reproduce within 6 h three key manifestations of end-stage Covid-19, we summarize here the unique features zymosan. It is extracted from the membrane of *Saccharomyces cerevisiae*, and consists of mannosylated cell wall proteins and highly branched β-glucans. The latter are D-glucose polymers linked by 1,3- and 1,6-β-glycosidic-bonds. Zymosan is widely used in immunological studies as a powerful stimulator of innate humoral and cellular immunity. The chemical structures (Fig. 8 a-d) tell little about zymosan’s real-life appearance (Fig. 8 e-i), i.e., 2-4 μm bean-shaped microparticles covered by knobs or bulges with extensions reminiscent of truncated bacterial pili.^45,46^

**Figure 8.**
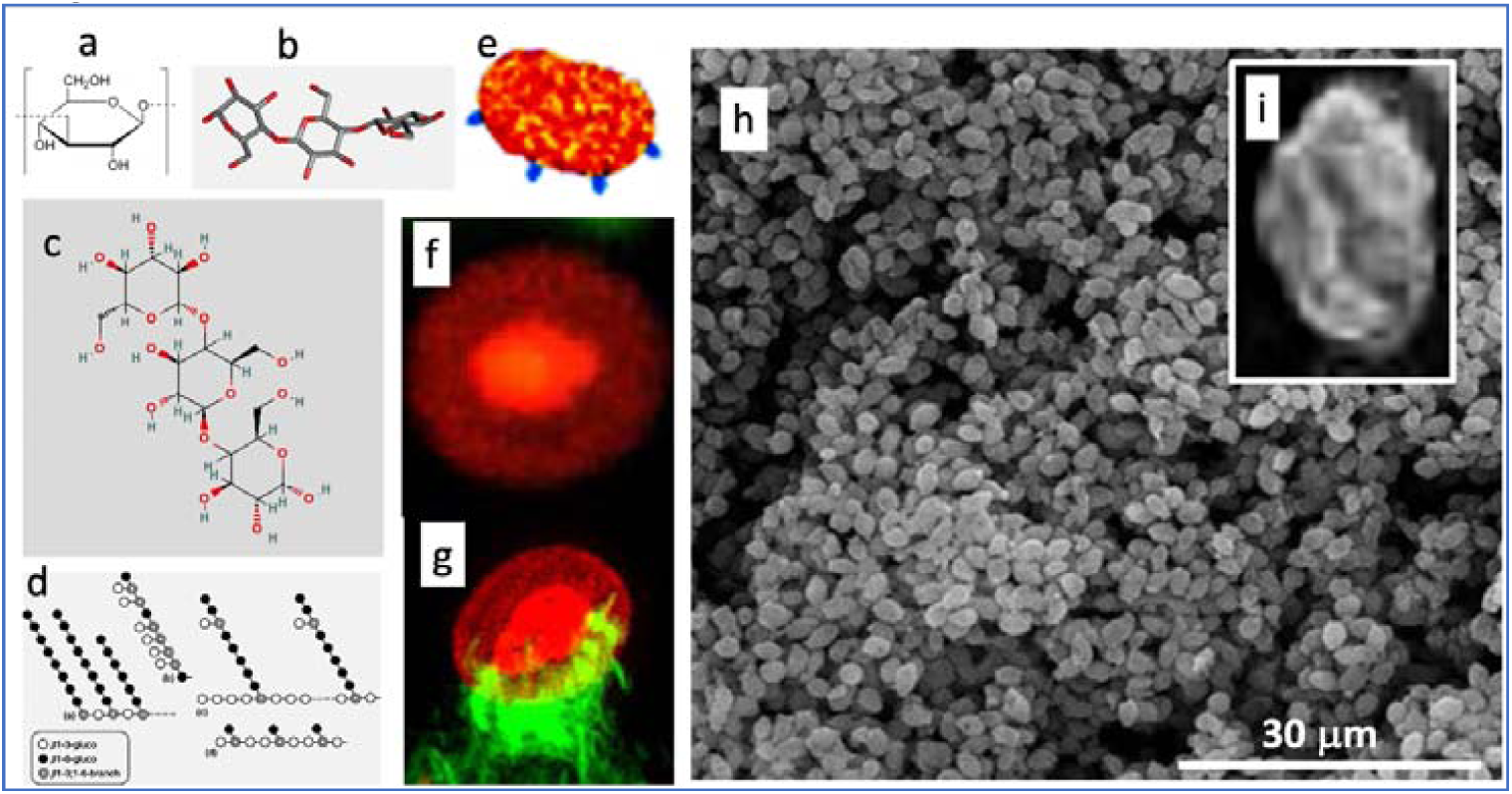
Various illustrations of zymosan. Panels A-D are chemical, graphical and 3D-steric presentation of structure, E is an artistic visualization and F-I are electron microscopic and scanning electron microscopic images of zymosan. The corona in F is with a fluorescent dye conjugated to zymosan, and G is a zymosan particle undergoing phagocytosis, where the green fluorescing “claws” are macrophage pseudopods embracing the particle as first step of phagocytosis. The insert I is a particle zoomed in from H. Free pictures from the internet and modified reproductions of figures in Ref. ^47^ with permission.

Zymosan was described as a potent activator of the C system 81 years ago,^48^ and has been used for modeling phagocytosis.^46^. Similarly to Toll-like receptor (TLR)-4 mediated LPS stimulation of NF-κB in immune cells, zymosan stimulates the production of inflammatory cytokines via Toll-like receptors TLR-2 and TLR-6.^49,50^ Furthermore, zymosan also activates another transmembrane signaling receptor, Dectin 1,^51-53^ which collaborates with TLR-2 in NF-κB mediated cytokine production.^54,55^. Due to all these redundant pro-inflammatory stimuli, zymosan has been used in many inflammatory disease models in mice and rats, as listed in Table 2.

**Table 2.**
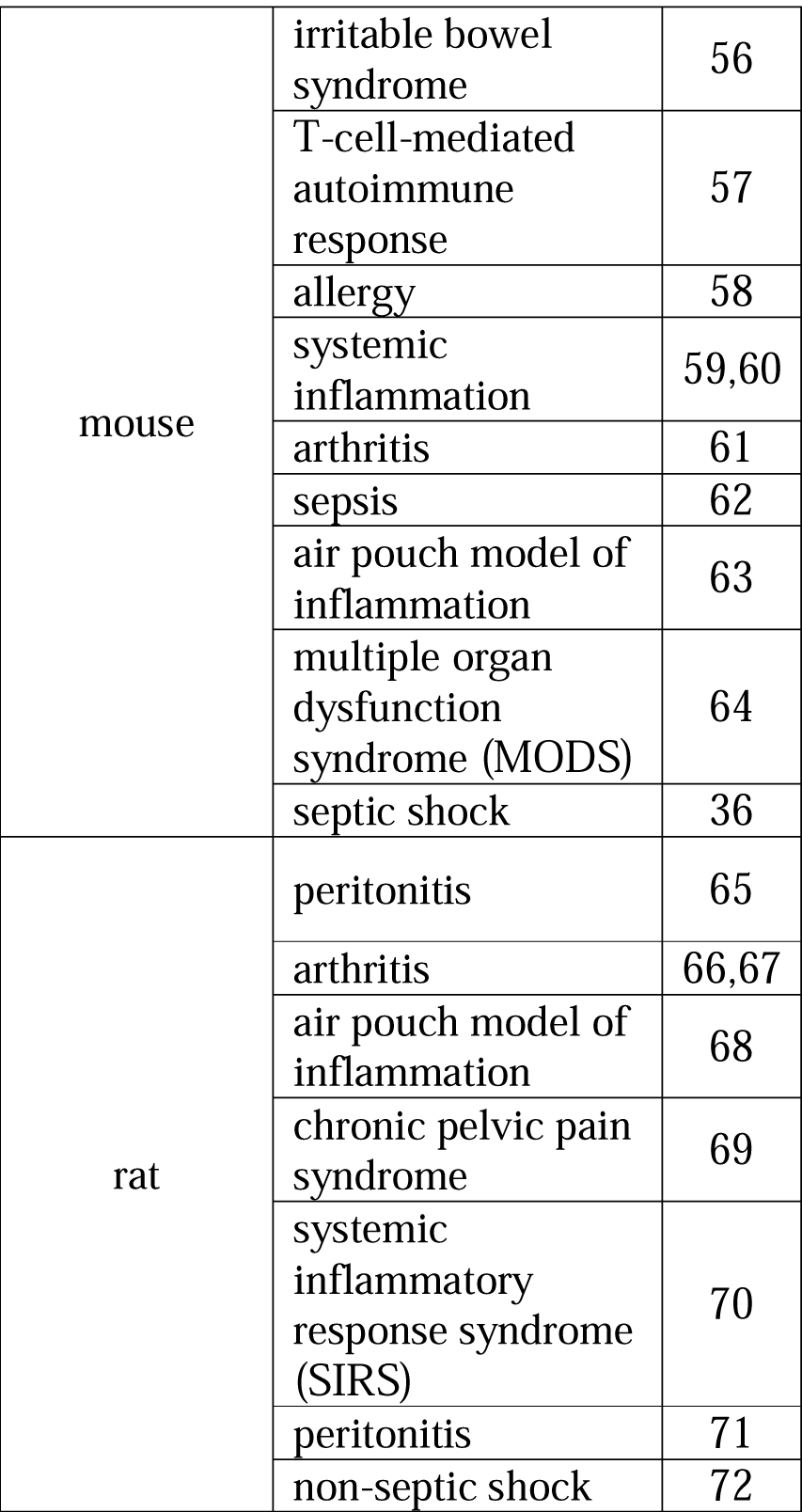
Murine models of inflammatory diseases using zymosan as inflammation inducer

|  |  |  |
| --- | --- | --- |
| mouse | irritable bowel syndrome | 56 |
|  | T-cell-mediated autoimmune response | 57 |
|  | allergy | 58 |
|  | systemic inflammation | 59,60 |
|  | arthritis | 61 |
|  | sepsis | 62 |
|  | air pouch model of inflammation | 63 |
|  | multiple organ dysfunction syndrome (MODS) | 64 |
|  | septic shock | 36 |
| rat | peritonitis | 65 |
|  | arthritis | 66,67 |
|  | air pouch model of inflammation | 68 |
|  | chronic pelvic pain syndrome | 69 |
|  | systemic inflammatory response syndrome (SIRS) | 70 |
|  | peritonitis | 71 |
|  | non-septic shock | 72 |

### Current animal models of CSS

Despite substantial efforts to develop effective pharmacotherapy against severe Covid-19, the standard of care today is based on traditional antibiotic and anti-inflammatory agents and some monoclonal antibodies, whose success is limited.^73-76^ One contributing reason for the shortage of new, more specific and effective drugs is the lack of appropriate, widely accessible animal model of Covid-19 or CSS. Natural and genetically modified species used to model different aspects of COVID-19 include mice, ferrets, cats, dogs, pigs, and non-human primates.^77-81^ The models described for CSS include the Staphylococcal superantigen mutant model in rabbits,^82^ the hemolytic transfusion model in mice,^83^ and the reactions of dogs to anti-CD28 mAb,^84^ or primates to simian immunodeficiency virus,^85^ or pigs to a virulent African swine fever virus.^86^. Yet another porcine model utilized LPS to induce CSS along with ARDS.^87^ However, none of these models can recapitulate the sustained immunopathology of patients with severe Covid-19 or CSS. Also, the use of gene-modified animals and high-containment BSL3+ facility in the case of infectious virus, are difficult to implement for high throughput drug testing in the pharmaceutical industry.

### Molecular and cellular mechanisms of zymosan’s induction in pigs of warning triad for bad prognosis in severe COVID-19

As to why it is possible in pigs to reproduce with zymosan 3 unfavorable disease markers in severe COVID-19, a likely answer is that zymosan is a very strong, multivalent stimulator of the innate immune system, a condition that also prevails in late stage COVID-19. Pigs, just as calves, sheep, goats and some other species, are very sensitive to innate immune stimulation.^88^ These species have pulmonary intravascular macrophages (PIM cells) in their lungs’ microcirculation which firmly attach to the capillary endothelium via junction-like intercellular adhesion plaques.^88^ These highly phagocytic and intense secretory macrophages with direct access to blood are also in close proximity to the smooth muscle cell layer of blood vessels, making these animals’ pulmonary arteries highly sensitive to the vasoactive mediators liberated upon encounter with, and phagocytic uptake of nanoparticles. These include TXA_2_, a strong vasoconstrictor eicosanoid, which is the prime suspect in the hemodynamic changes caused by iv. nanoparticles in pigs.^88^ Zymosan can stimulate these macrophages via 3 independent pathways; one via the anaphylatoxin (C3a, C5a) receptors, another is via TLR-2/6, and the third one is the Dectin-1 receptors.^88^ These redundant activation pathways explain the high inter-animal reproducibility of hemodynamic changes caused by zymosan. That zymosan exerts its vasoactivity and pro-inflammatory cytokine inducing effects via cells exposed to plasma is supported by the finding that the kinetics of liberation of TXB_2_ and inflammatory cytokines in an *in vitro* peripheral blood mononuclear cell model of CSS^38^ is very similar to those seen in pigs, namely immediate production of TXB_2_ and slower release of cytokines on a time scale of hours.^38^

Regarding the lymphopenia and its correlation with IL-6 gene expression (Fig. S1), IL-6 is known to upregulate the pro-apoptotic Fas, resulting in the loss of mature lymphocytes.^89^ High levels of IL-6 might also reduce lymphocyte count by inhibition of lymphopoiesis in the bone marrow.^90^ The neutrophil granulocytosis, in turn, is a common sign of strong inflammation with cytokine release, a well-known disease marker. As for the roles of CCL2 (*C-C motif chemokine ligand 2*, also known as monocyte chemoattractant protein 1, MCP1) and CXCL (*C-X-C motif chemokine ligand 10*, also known as interferon gamma-induced protein 10, IP-10), we have no information in the literature that would suggest a direct role of these chemokines in rising the NLR.

### The utility of porcine zymosan-induced CSS model

The experimental procedures applied in this study represent relatively straightforward in vivo investigation of systemic flare-up of inflammatory processes in the body, a complex immune phenomenon, a feared end-stage of many severe diseases including COVID-19, viral infections,^91^ monoclonal antibody and CAR-T-cell therapies,^92,93^ acute respiratory disease syndrome^94^ and multiorgan failure.^94,95^ CSS has multiple manifestations, and the different models discussed above focus on different endpoints. In the present model we have focused on three standard physiological parameters which have been reported as bad prognostic indicators in late-stage COVID-19; pulmonary hypertension, rise of NLR and cytokine release, which can be also common features of all CSS, regardless of cause. This choice of endpoints, taken together with the increasing appreciation of pigs, as an immune toxicology model,^34^ the inexpensive access to zymosan, the rapid (up to 6 h) experimentation, the avoidance of problematic interpretation of immune data in murine models, exotic animals or infectious viruses with need for BSL-3 facility, or sophisticated gene technology, suggests that the porcine zymosan-induced CSS model may provide a new tool to better understand and develop effective pharmacotherapies against CSS in general, and end-stage COVID-19, in particular. Although the hemodynamic data were critical in realizing the relevance of the model for end-stage COVID-19, they are not essential as disease markers, implying that performing only blood WBC counting may serve the purpose of streamlined drug screening, thus advancing progress in preventing or attenuating CSS.

## Acknowledgements

The expert technical supports by Katalin Simay, Henriett Biró and Krisztina Fazekas are gratefully acknowledged.

